# Regulation of angiotensin converting enzyme 2 (ACE2) in obesity: implications for COVID-19

**DOI:** 10.1101/2020.04.17.046938

**Authors:** Saba Al Heialy, Mahmood Hachim, Abiola Senok, Ahmad Abou Tayoun, Rifat Hamoudi, Alawi Alsheikh-Ali, Qutayba Hamid

**Author notes:** Corresponding author: Professor Qutayba Hamid.

## Abstract

The ongoing COVID-19 pandemic is caused by the novel coronavirus SARS-CoV-2. Age, smoking, obesity, and chronic diseases such as cardiovascular disease and diabetes have been described as risk factors for severe complications and mortality in COVID-19. Obesity and diabetes are usually associated with dysregulated lipid synthesis and clearance which can initiate or aggravate pulmonary inflammation and injury. It has been shown that for viral entry into the host cell, SARS-CoV-2 utilizes the angiotensin converting enzyme 2 (ACE2) receptors present on the cells. We aimed to characterize how SARS-CoV-2 dysregulates lipid metabolism pathways in the host and the effect of dysregulated lipogenesis on the regulation of ACE2, specifically in obesity. In our study, through the re-analysis of publicly available transcriptomic data, we first found that lung epithelial cells infected with SARS-CoV-2 showed upregulation of genes associated with lipid metabolism, including the *SOC3* gene which is involved in regulation of inflammation and inhibition of leptin signaling. This is of interest as viruses may hijack host lipid metabolism to allow completion of their viral replication cycles. Furthermore, a mouse model of diet-induced obesity showed a significant increase in *Ace2* expression in the lungs which negatively correlated with the expression of genes that code for sterol response element binding proteins 1 and 2 (SREBP). Suppression of *Srebp1* showed a significant increase in *Ace2* expression in the lung. Together our results suggest that the dysregulated lipogenesis and the subsequently high ACE2 expression in obese patients might be the mechanism underlying the increased risk for severe complications in those patients when infected by SARS-CoV-2.

## Introduction

As the COVID-19 pandemic is accelerating, people worldwide are warned to take necessary precautions to avoid infection. With changing statistics every day, it is clear that certain groups of individuals are at increased risk of severe infection. In particular, the groups at risk are the elderly, individuals with chronic health conditions such as diabetes, cancer and cardiovascular diseases. Most cases of COVID-19 are classified as mild or moderate. However, 14% of the cases are severe and may lead to acute respiratory distress syndrome (ARDS) and even death in those infected (1). Therefore, it is crucial to understand the mechanism by which this virus causes organ injury and, in particular the immune system which is mounted in response to the infection. A few studies have now identified other risk factors for severe complications and death due to COVID-19, namely smoking and obesity (2, 3). The findings related to obesity are plausible since obese individuals tend to be more difficult to intubate, and excess body weight may contribute to increased pressure on the diaphragm which may make breathing more difficult during infection. Moreover, it is well established that obesity leads to chronic meta-inflammation even in the absence of infection which has detrimental effects on the immune system (4). In addition, obesity is associated with dysregulated lipid synthesis and clearance which can initiate or aggravate pulmonary inflammation. It has also been shown that antiviral medication and vaccines are less effective in obese individuals (5). In relation to the influenza virus, obesity may have a role in the viral life cycle which along with a dysregulated immune system could lead to severe complications (6). During the H1N1 pandemic in 2009, obesity was classified as an independent risk factor for hospitalization, need for mechanical ventilation and death. These observations are concerning since over one third of the world population are classified as overweight or obese (7). Therefore, as this current COVID-19 pandemic is increasing, it is important to understand the molecular mechanisms through which obesity increases the complications related to COVID-19 to hopefully be able to design more appropriate therapies. Moreover, understanding the effects of obesity on COVID-19 may shed light on the pathogenecity of SARS-CoV-2.

The mode of cellular entry of the novel severe acute respiratory syndrome coronavirus, SARS-CoV-2, is through its binding to the angiotensin converting enzyme 2 (ACE2) and is similar to SARS-CoV responsible for the 2003 pandemic (8). Specifically, the spike glycoprotein on the virion binds to peptidase domain of ACE2. Physiologically, ACE2 is part of the renin-angiotensin system (RAS) and serves as a key regulator of systemic blood pressure through the cleavage of Angiotensin (Ang) I to generate the inactive Ang 1-9 peptide and it directly metabolizes Ang II to generate Ang 1-7 limiting its effects on vasoconstriction and fibrosis. Other than serving as a functional receptor for SARS-CoV, ACE2 has been shown to be implicated in cardiovascular pathologies, diabetes and lung disease. It is expressed by cells of the heart, kidney and more specifically in lung epithelial cells (9). Although ACE2 expression correlates with susceptibility of SARS-CoV infection, the relationship between ACE2 and SARS-CoV-2 remains unclear. In fact, studies have suggested a protective role for ACE2 where overexpression of ACE2 attenuates lung inflammation (10). Current research is focusing on the regulation and role of this receptor in relation to SARS-CoV-2. The aim of this study was to identify mechanisms, through re-analysis of publicly available transcriptomic data, by which SARS-CoV-2 dysregulates the lipid mechanism pathways and investigate the effect of dysregulated lipogenesis on the regulation of ACE2, specifically in obesity.

## Materials and Methods

### Differentially expressed genes in bronchial epithelial cells infected with SARS-CoV-2

In order to identify essential differentially expressed genes in SARS-CoV-2 infected versus non-infected epithelial cells, we re-analyzed the publicly available transcriptomic dataset (GSE147507) recently uploaded to the Gene Expression Omnibus (GEO) (11). Independent biological triplicates of primary human lung epithelium (NHBE) were mock-treated or infected with SARS-CoV-2 (USA-WA1/2020), then subjected to whole transcriptomic analysis using RNA-Sequencing on Illumina Next Seq 500. The Raw Read Counts were retrieved and filtered from non-expressing genes that showed zero counts in the six samples. Out of the original 23710 genes, only 15487 were expressing and selected for further analysis. The filtered gene expression was uploaded to AltAnalyze software for Comprehensive Transcriptome Analysis(12). Principle component analysis and Heatmap clustering were generated, and differentially expressed genes (DEGs) were identified using LIMMA algorithm built in AltAnalyze software. The genes that made the optimal hierarchal cosine clustering were identified, and the common pathways shared by these genes are listed according to their significance. Graphical visualization of the gene was made using the Metascape online tool for gene ontology (http://metascape.org) (13).

## Results

### SARS-CoV-2 differentially expressed genes related to lipid metabolism in epithelial cells

121 DEGs in infected versus non-infected cells were identified, and they clustered the two groups separately (Figure 1). As expected, the top pathways where the DEGs by SARS-CoV-2 are involved were related to inflammatory, immune, cytokines, and antiviral responses.

**Figure 1:**
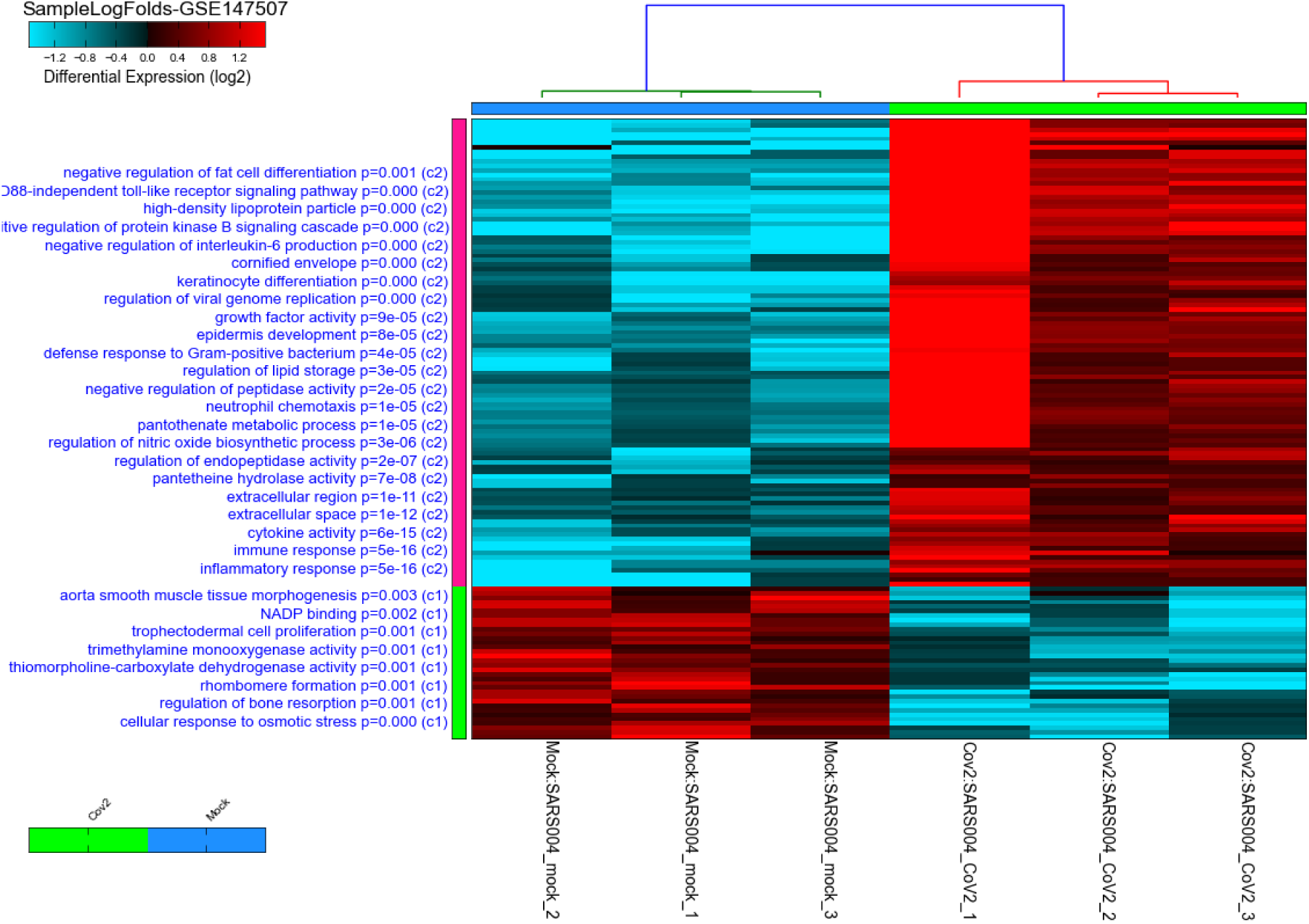
Heatmap and clustering of SARS-CoV-2 infected healthy epithelial cells versus noninfected (mock-infected cells) using GSE147507 publicly available transcriptomics dataset. Pathways, where the DEGs are involved, are shown on the right side of the heatmap with their adjusted p-value. The C1 represent pathways of genes downregulated in infected cells, while C2 represent pathways of genes upregulated when cells are infected compared to mock-infected cells.

### Top DEGs in infected cells can have a role in white fat differentiation

Of note, as shown in Figure 1, genes involved in lipid storage and high-density lipoprotein particles were among the top upregulated genes by the virus. To have a detailed analysis of the pathways where the top DEGs in infected versus non-infected epithelial cells are involved, we uploaded the 121 DEGs to metascape online tool. Again, most of the DEGs were involved in immune response-related such as Interleukin (IL)-17 signaling pathway. This result was expected and validated our bioinformatics analysis (14). IL-10 signaling pathway, acute inflammatory response, metal sequestration by antimicrobial proteins, defense response to another organism, negative regulation of apoptotic signaling pathway, acute-phase response, cellular response to tumor necrosis factor, response to antibiotic and modulation by a host of the viral process (Figure 2 and table 1) were also involved. Other sets of enriched pathways are related to cell and tissue homeostasis like positive regulation of cell migration, blood vessel morphogenesis, regulation of bone resorption, activation of matrix metalloproteinases, and regulation of smooth muscle cell proliferation. The third set was related to metabolic pathways like negative regulation of ion transport, regulation of glucose metabolic process, the release of cytochrome c from mitochondria, and regulation of fat cell differentiation.

**Figure 2:**
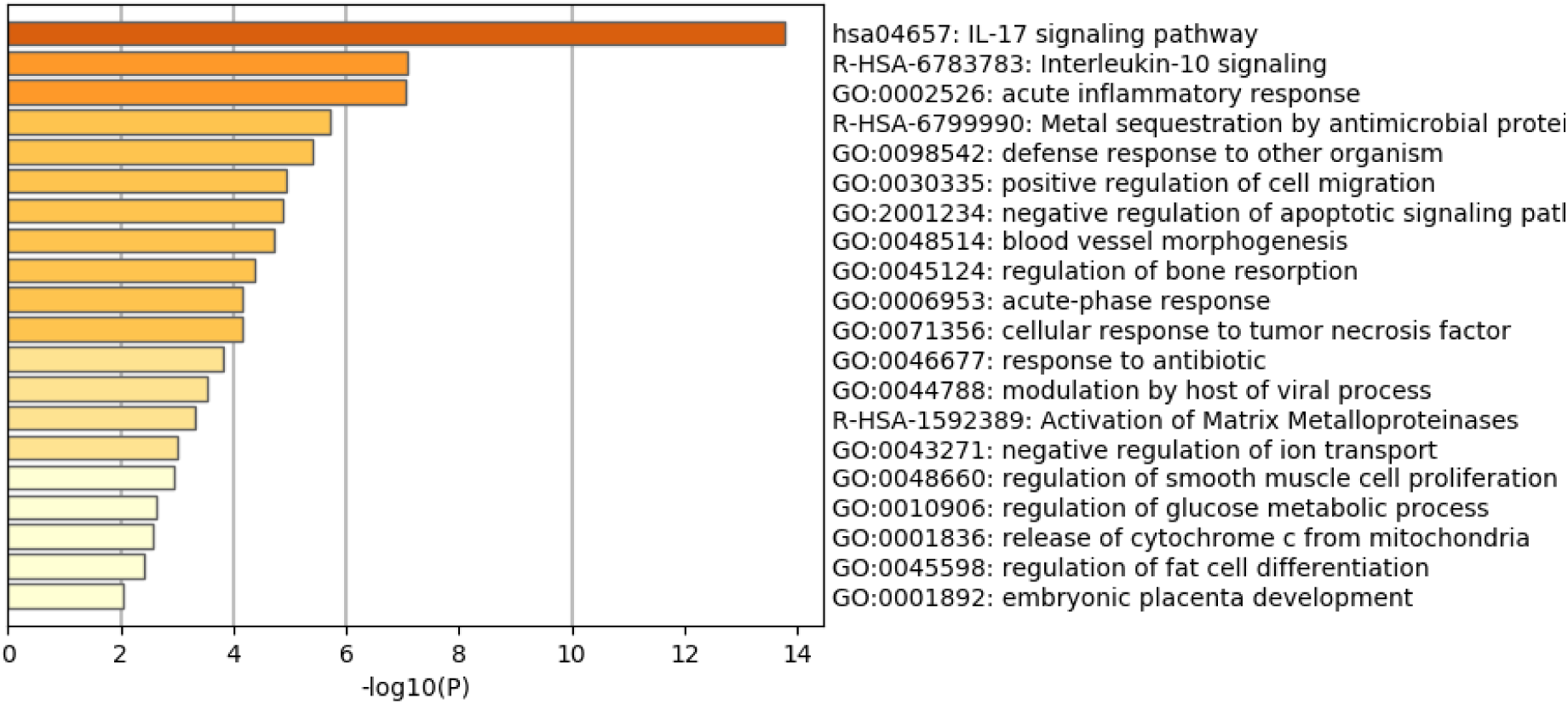
Heatmap of top Gene Ontology (GO) of the top DEGs in infected normal epithelium versus mock-infected cells. The bars represent the -log10 of the adjusted p-value for each pathway.

**Table 1:** Top Gene Ontology (GO) of the top DEGs in infected normal epithelium versus mock-infected cells and the symbols of genes in each pathway

| Category | Term | Description | LogP | Log(q-value) | InTerm_InList | Symbols |
| --- | --- | --- | --- | --- | --- | --- |
| KEGG Pathway | hsa04657 | IL-17 signaling pathway | -13.7848 | -9.466 | 12/93 | *CSF2, CSF3, CXCL3, IL6, CXCL8, MMP9, MMP13, S100A7, S100A8, CCL20, CXCL5, TNFAIP3, C3, CAMP, ICAM1, MAOB, IL36G, PGLYRP4, ZC3H12A, KLHL6, GSDMA, SSC5D, SERPINA3, LTB, MMP11, TNFSF14, SEMA3G, C1QTNF1, ADAM32, S100A7A, CSF1R, IFI6, IFI27, MX1, SAA1, SOD2, XAF1, CFB, C8B, SAA4, EGR2, ESRRB, PDK4* |
| Reactome Gene Sets | R-HSA-6783783 | Interleukin-10 signaling | -7.09786 | -3.779 | 6/47 | *CSF2, CSF3, ICAM1, IL6, CXCL8, CCL20, LTB, CXCL5, C3, CXCL3, MMP9, SAA1, C8B, CAMP, TNFAIP3, EGR2, CSF1R, MX1* |
| GO Biological Processes | GO:0002526 | acute inflammatory response | -7.05687 | -3.779 | 10/222 | *SERPINA3, CFB, C3, C8B, ICAM1, IL6, S100A8, SAA1, SAA4, VNN1, MMP9, TNFAIP3, ZC3H12A, IL36G, CAMP, PGLYRP4* |
| Reactome Gene Sets | R-HSA-6799990 | Metal sequestration by antimicrobial proteins | -5.71647 | -2.740 | 3/6 | *S100A7, S100A8, S100A7A, TNFAIP3, VNN1, CAMP, PGLYRP4, SERPINA3, C3, CSF2, IL6, CXCL8, MMP9, ZC3H12A, ICAM1, TCIM, SOD2, RNF183, IFI6* |
| GO Biological Processes | GO:0098542 | defense response to other organism | -5.39645 | -2.446 | 13/596 | *CAMP, IFI6, IFI27, IL6, MX1, S100A7, S100A8, CCL20, TNFAIP3, PGLYRP4, ZC3H12A, GSDMA, SSC5D* |
| GO Biological Processes | GO:0030335 | positive regulation of cell migration | -4.93817 | -2.096 | 12/560 | *CSF1R, ICAM1, IL6, CXCL8, MMP9, MYLK, S100A7, CCL20, SOD2, TNFSF14, SEMA3G, ZC3H12A, SAA1, C3, S100A8, C1QTNF1, MIR503* |
| GO Biological Processes | GO:2001234 | negative regulation of apoptotic signaling pathway | -4.87469 | -2.047 | 8/233 | *CSF2, IFI6, ICAM1, MMP9, SOD2, TNFAIP3, VNN1, PAK5, S100A8, RNF183, IFI27* |
| GO Biological Processes | GO:0048514 | blood vessel morphogenesis | -4.719 | -1.931 | 13/690 | *C3, IL6, CXCL8, MYLK, S100A7, TNFAIP2, TNFAIP3, FOXN1, KLF2, VASH1, ZC3H12A, THSD7A, MIR503, SOD2, MMP9, VNN1, C8B, ICAM1, KLHL6, SAA1, CSF1R, CSF2, LTB, SSC5D, C1QTNF1* |
| GO Biological Processes | GO:0045124 | regulation of bone resorption | -4.3687 | -1.652 | 4/42 | *CSF1R, IL6, PDK4, TNFAIP3, SERPINA3, ESRRB, CSF2, CSF3, MMP9, ICAM1, FOXN1, VNN1, ZC3H12A, S100A8* |
| GO Biological Processes | GO:0006953 | acute-phase response | -4.17408 | -1.492 | 4/47 | *SERPINA3, IL6, SAA1, SAA4, CXCL8* |
| GO Biological Processes | GO:0071356 | cellular response to tumor necrosis factor | - 4.16725 | -1.492 | 8/293 | *ICAM1, CXCL8, LTB, CCL20, TNFAIP3, TNFSF14, KLF2, ZC3H12A, IL6, FOXN1* |
| GO Biological Processes | GO:0046677 | response to antibiotic | -3.8017 | -1.253 | 8/331 | *CSF3, ICAM1, IL6, MAOB, S100A8, TNFAIP3, KLF2, ZC3H12A, CXCL8, CCL20, CSF2, MMP9, SOD2, IFI6, PDK4, NID1, SAA1, TNFSF14, VNN1, C1QTNF1, MIR503, PFKFB1* |
| GO Biological Processes | GO:0044788 | modulation by host of viral process | - 3.53447 | -1.055 | 3/28 | *CSF1R, IFI27, ZC3H12A, IFI6, MX1, XAF1, CAMP, PGLYRP4, IL6, TNFAIP3, CXCL8, ICAM1* |
| Reactome Gene Sets | R-HSA-1592389 | Activation of Matrix Metalloproteinases | - 3.32026 | -0.899 | 3/33 | *MMP9, MMP11, MMP13, IL6, ICAM1, NID1, COLGALT2, EGR2, PDK4, PFKFB1, TNFAIP3, KLF2, SERPINA3, ADAM32* |
| GO Biological Processes | GO:0043271 | negative regulation of ion transport | - 3.01754 | -0.715 | 5/162 | *ICAM1, MAOB, MMP9, KCNE3, KCNRG, TNFAIP3, ZC3H12A, SSC5D* |
| GO Biological Processes | GO:0048660 | regulation of smooth muscle cell proliferation | - 2.93566 | -0.673 | 5/169 | *IL6, MMP9, SOD2, TNFAIP3, MIR503* |
| GO Biological Processes | GO:0010906 | regulation of glucose metabolic process | - 2.63653 | -0.448 | 4/119 | *ESRRB, PDK4, PFKFB1, C1QTNF1* |
| GO Biological Processes | GO:0001836 | release of cytochrome c from mitochondria | - 2.58309 | -0.422 | 3/59 | *IFI6, MMP9, SOD2, SERPINA3, C3, TNFSF14* |
| GO Biological Processes | GO:0045598 | regulation of fat cell differentiation | - 2.40524 | -0.306 | 4/138 | *IL6, MMP11, ZC3H12A, PTPRQ, EGR2* |
| GO Biological Processes | GO:0001892 | embryonic placenta development | - 2.05701 | -0.084 | 3/91 | *CSF2, ESRRB, VASH1* |

De novo cellular lipogenesis, if disturbed, can change cell deformability as it influences the phospholipid composition of cellular membranes and, as a consequence, can disturb transmembrane receptors like growth factor receptor needed for cell survival (15). Based on that we were interested in deciphering the effect of viral infection on epithelial cells lipid metabolism pathways and how the disturbed lipid metabolism, like in obesity and diabetes, might worsen the condition of COVID-19

### Negative regulator of lipogenesis was downregulated in infected cells

In this study, *IL6, MMP11, ZC3H12A, PTPRQ*, and *EGR2* were found to share a common pathway related to the regulation of fat cell differentiation. Of interest, *PTPRQ* and *EGR2* showed significant downregulation in infected cells compared to mock-infected epithelial cells, as shown in Figure 3. PTPRQ is protein tyrosine phosphatases (PTPs) that regulate tyrosine phosphorylation in signal transduction, and the encoded protein is a negative regulator of mesenchymal stem cell differentiation into adipocytes (16). It is synthesized in the lung and kidney and is downregulated in the early stages of adipogenesis(17). Recent studies linked downregulated PTPs like PTPRQ to lower weight gain, food intake, and leptin resistance(18).

**Figure 3:**
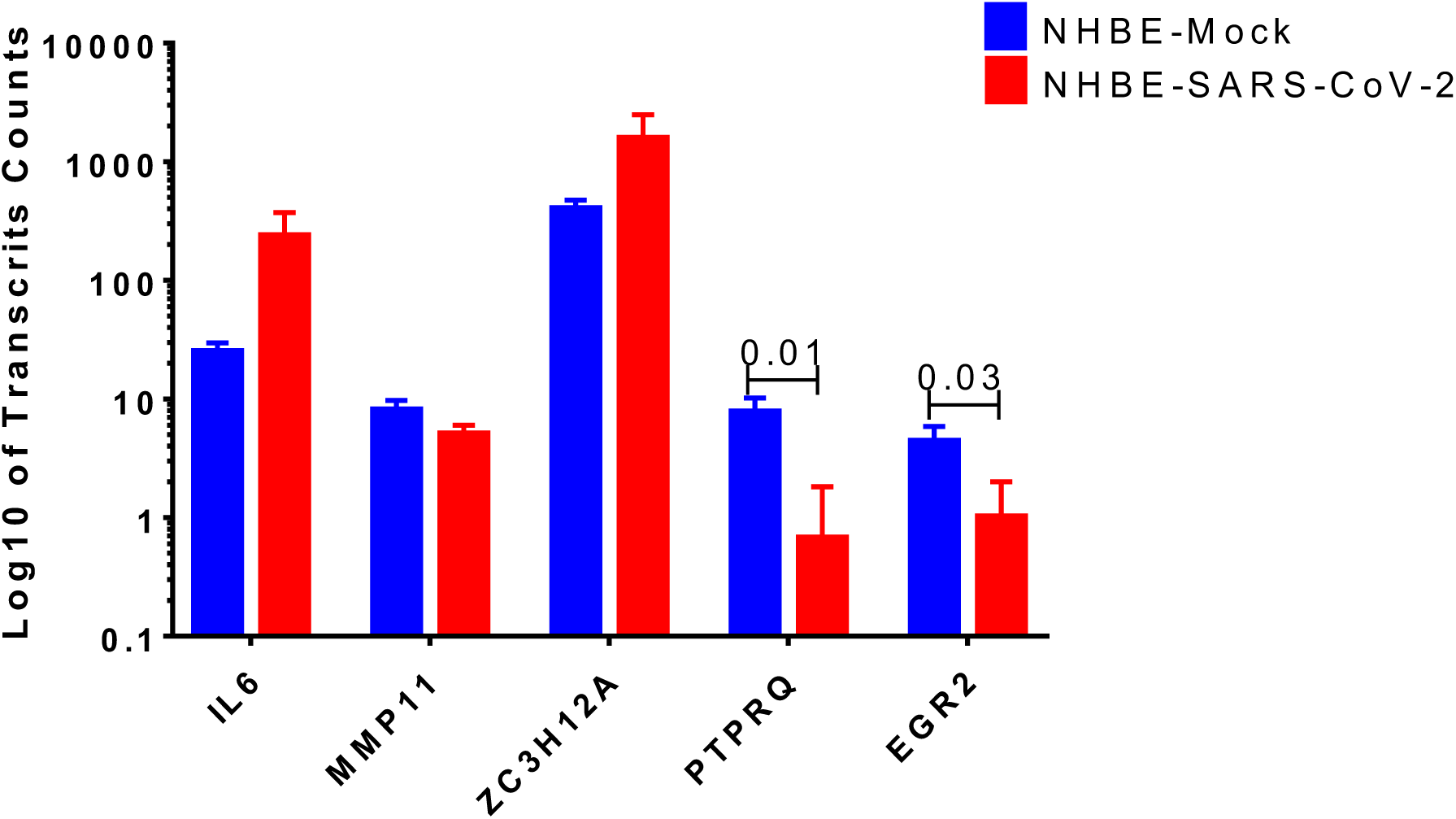
log10 of transcripts counts of the genes related to the regulation of fat cell differentiation *IL6, MMP11, ZC3H12A, PTPRQ*, and *EGR2* in infected cells compared to mock-infected epithelial cells.

### SARS-CoV-2 upregulate leptin signaling regulator SOCS-3

Next, we tried to look for the trend of changes in the expression of genes involved in lipid metabolism (although the differences were not statistically significant) but this data can give us an idea about the deranged lipid-related pathways. To visualize the leptin signaling pathways, the filtered gene expression was uploaded to PathVisio pathway analysis and drawing software(19). As shown in Figure 4, there is derangement of a leptin signaling pathway in terms of upregulation or downregulation of genes involved, of note the *SOCS3, STAT1*, *NFKB1* and *IL1B* were the top upregulated genes. Most individuals with obesity have leptin resistance by leptin and its receptor inhibitor *SOCS-3* (suppressor of cytokine signaling-3), leading to dysfunction of leptin biological function(20)

**Figure 4:**
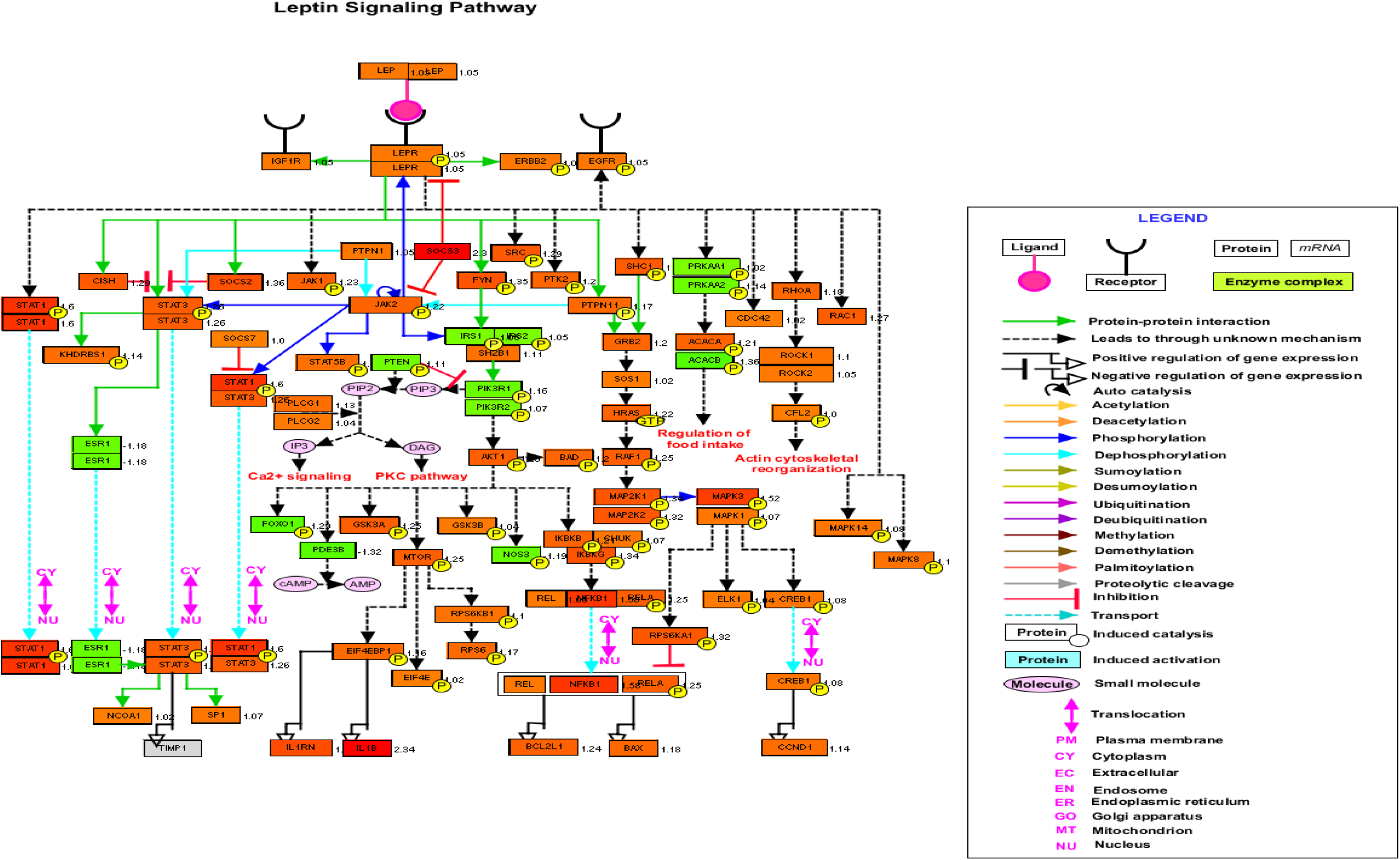
Leptin signaling pathways genes members and their expression in SARS-CoV-2 infected epithelial cells versus noninfected. Red color means upregulation in infected cells, and green means downregulated as generated by PathVisio software.

### Lung ACE2 expression was significantly upregulated in obese mice

We then questioned whether disturbed lipid metabolism in obesity could affect the major players’ host genes involved in virus binding and entry, namely ACE2. Apart from the circulating RAS, the local lung-based RAS plays a specific role in the injury/repair response(21) and recently was documented to have a pro-fibrotic effect independent of the known blood pressure effect(22). Recently, a note that COVID-19 can induce RAS imbalance that drives acute lung injury(23). *In vitro* results showed that continued viral infection would reduce membrane ACE2 expression, leading to unstoppable activation of RAS in the lungs, which further induce local inflammation by recruited neutrophils after LPS stimuli(24). In COVID-19, ACE2 showed opposite harmful effects as an entry point, and beneficial effect by counteracting the overstimulated RAS as it degrades AngII to angiotensin 1-7 (Ang1-7) (25). Interestingly Ang1-7 is shown to block high-fat diet-induced obesity, which increased ACE2 expression in adipose tissue(26). Therefore, our hypothesis was that induced obesity can upregulate ACE2 in the lung in response to a high-fat diet, which makes the lung more susceptible to viral entry but can regulate the overstimulated RAS. To examine that, we explored publicly available transcriptomics data of where lungs were examined after inducing obesity to look for the ACE2 expression changes. GSE38092 dataset was found with eight regular weight mice versus eight diet-induced obese mice where their lungs were extracted for microarray gene expression profiling(27), as shown in Figure 5C. Interestingly, lung *Ace2* expression was significantly upregulated in obese mice compared to lean.

**Figure 5:**
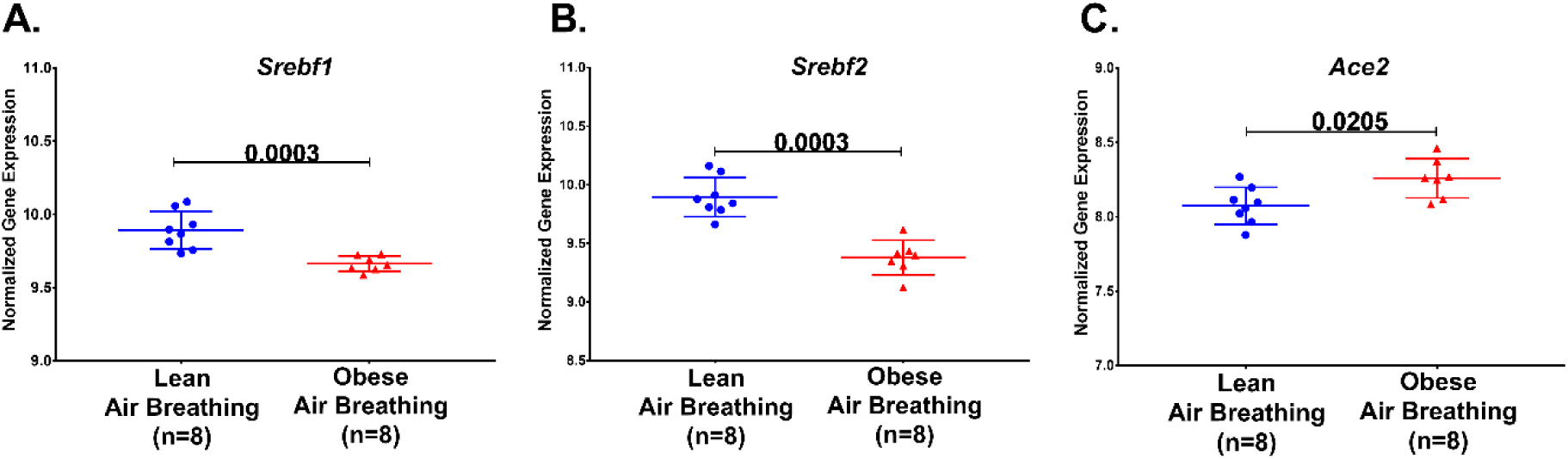
*Srebf1*, *Srebf2* and *Ace2* mRNA normalized gene expression in response to a high-fat diet in obese compared to regular weight mice extracted from publicly available transcriptomic dataset GSE38092

Sterol-response element binding proteins (SREBP) are transcription factors that have been associated with lipogenesis, adipogenesis and cholesterol homeostasis to prevent lipotoxicity. Studies have shown differential expression of SREBP-1 in regard to obesity. Figure 5 A and B shows a decrease in *Srebf1* and *Srebf2*, the genes that code for the different proteins, namely SREBP-1 and SREBP-2.

The increased level of *Ace2* in the lungs of obese mice led us to investigate which cell type in the lung has the highest expression of *Ace2*. We explored LungGENS web-based tool that can map single-cell gene expression in lung(28). Among all cells in the human lungs, *Ace2* was expressed exclusively by epithelial cells, as shown in Figure 5.

### Lung epithelial cells express lesser ACE2 during activated lipogenesis

The next question is how diet-induced obesity mechanistically can upregulate ACE2 in the lung, to answer this question, another dataset (GSE31797) was explored where the dynamics of lung lipotoxicity was examined by manipulating SREBP (29). SREBPs regulate the expression of genes involved in lipid synthesis and function by their actions as transcription factors(30). It was shown previously that the deletion of ACE2 in the liver and skeletal muscles could induce lipogenesis by inducing SREBPs, indicating the role of the ACE2/Ang1-7 axis in lipid metabolism (31, 32).

**Figure 6:**
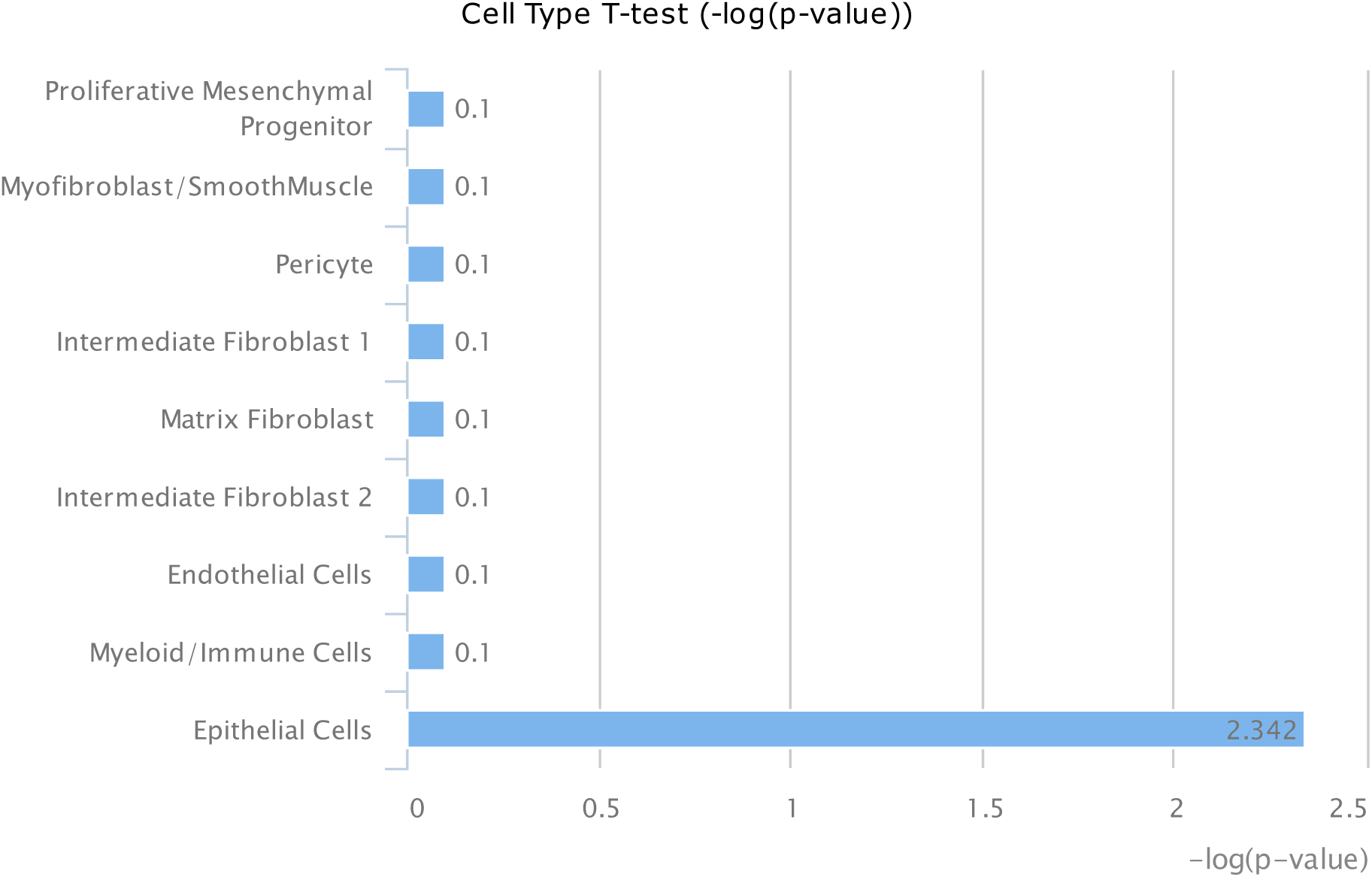
Lung cells type ACE2 mRNA expression using LungGENS web-based tool

In this dataset, alveolar type 2 cell RNA from Insig1/2Δ/Δ (activated SREBP1 levels) and Insig1flox/flox/Insig2−/− (suppressed SREBP1 levels) mice were profiled by microarray. Interestingly, activating SREBP1, coded by *Srebf1*, downregulated the expression of the three probes of the *Ace2* gene used in the microarray, as shown in Figure 7. This data indicates that *Ace2* expression might be under the control of SREBP1.

**Figure 7:**
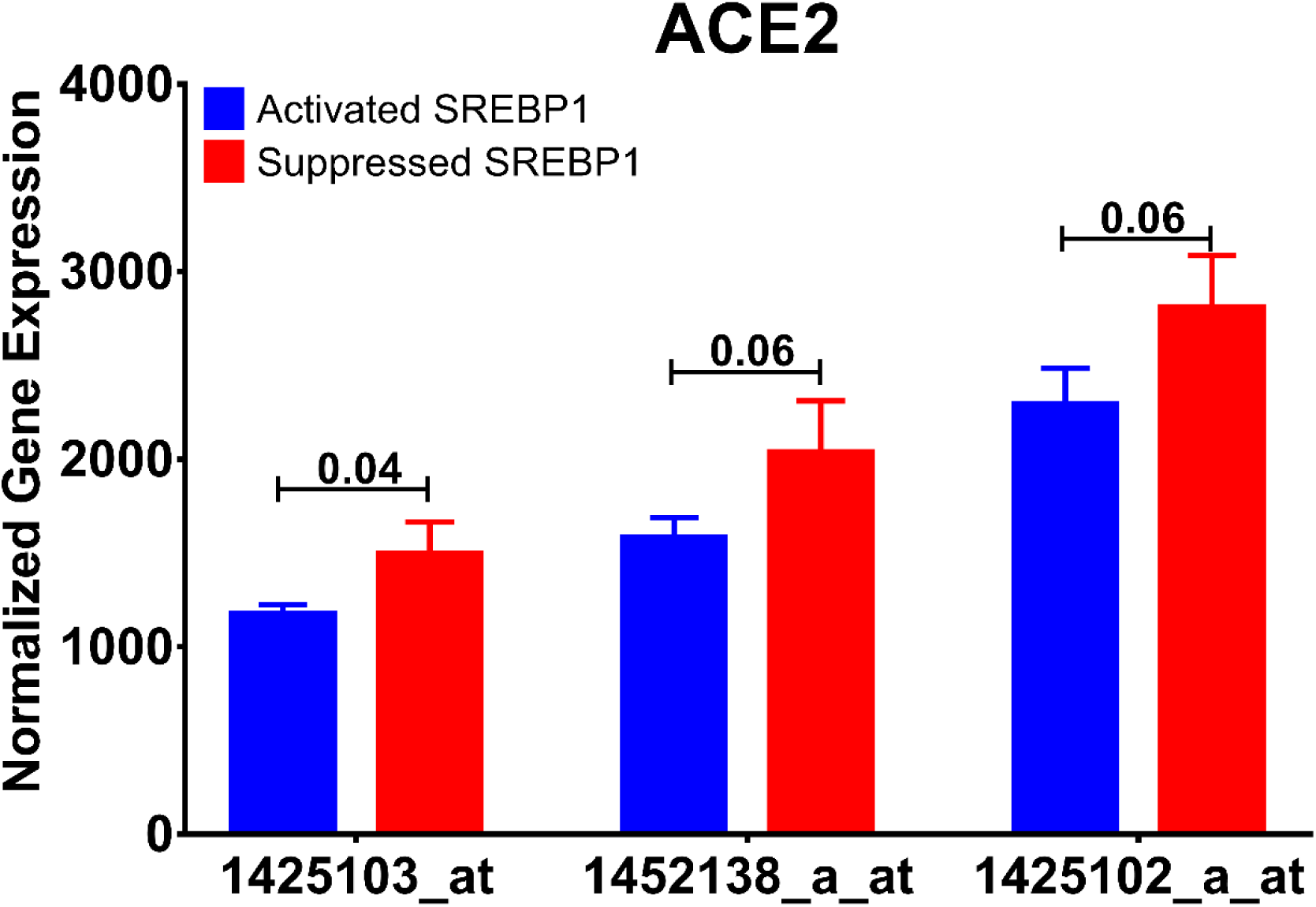
Normalized mRNA expression of Ace2 gene probes used in the publicly available dataset (GSE31797) comparing SREBP activated with SREBP inhibited alveolar cells.

## Discussion

This study, which uses publicly available data, demonstrates that firstly, SARS-CoV-2 infection induces changes in lipid profile in healthy hosts as demonstrated in infection of healthy epithelial cells. Secondly, this is important when obesity is at play as the lipid profile is already disrupted which may lead to increased susceptibility to infection which if occurs will further alter the lipid profile inducing hyper-inflammation.

We first demonstrated that SARS-CoV-2 infection of healthy epithelial cells compared to mock-infected cells clusters the genes involved in inflammatory, immune and viral responses (Fig. 1). In particular, the IL-17 and IL-10 signalling pathways were heavily impacted. Symptoms of severe COVID-19 have been associated with a cytokine storm with high levels of IL-17, IL-10, IL-1β, IL-2, IL-7, IL-8, IL-9 among many other pro-inflammatory cytokines (14, 33). IL-17, with its many pro-inflammatory effects, has been suggested as a potential target for the treatment of COVID-19. This is of interest as obesity is associated with high levels of immune cells producing IL-17 (34). IL-10, an anti-inflammatory cytokine with antiviral properties, is usually downregulated in infections. However, severe cases of COVID-19 have been associated with high levels of IL-10.

Obesity is a major health problem associated with increased risk of developing diabetes, hypercholesteremia and hypertension. Alterations in the metabolic pathways are seen such as increases in leptin and insulin secretion and decrease in adiponectin. Leptin stimulates fatty acid oxidation and may lead to lipotoxicity through decreased lipid accumulation in non-adipose tissue. As metabolic regulation and immune responses seem to be integrated with the function of one being dependent on the other, obesity is associated with high levels of pro-inflammatory mediators (35). Therefore, we focused our study on metabolic pathways in conjunction with inflammatory pathways. Our data revealed that among the pathways that were differentially expressed were the regulation of glucose metabolic process and regulation of fat cell differentiation (Fig. 2). Firstly, we found that *PTPRQ* and *EGR2* genes were significantly downregulated in SARS-CoV-2 infected healthy epithelial cells. Although they have been described to have roles in lipogenesis, their exact role in viral infection remains unknown and warrants further investigation.

The upregulation of *LEPR* and *LEP* and associated *SOC3* in response to SARS-CoV-2 infection was in line with mechansims which are dysregulated in the state of obesity. This finding is of interest as it has been previously shown that viruses such as the West Nile Virus hijack cellular cholesterol to redistribute it and allow completion of its replication cycle (36, 37). Previous studies have shown that obesity is associated with leptin resistance and increased blood levels of leptin with concomitant increases in *SOC3* which plays a role in inhibiting signal transduction of leptin and other cytokines (38).

Having established that SARS-CoV-2 infection and obesity share common pathways associated with dysregulation of lipid metabolism, we were interested to see if obesity, which has been described as a risk factor of COVID-19, is associated with higher susceptibility to infection. SARS-CoV-2 uses the ACE2 as a receptor for viral entry so we hypothesized that obesity may lead to higher expression of ACE2. Using a high-fat diet animal model of obesity, our results revealed a higher expression of *Ace2* among diet-induced obese mice compared to lean mice. The data also suggests that the expression of ACE2 is exclusive to lung epithelial cells. Previous studies with SARS-CoV have shown that infection state correlates with state of cell differentiation and expression of *ACE2* (39).

The present study focused on the relationship between ACE2 expression and dysregulation of lipid metabolism. Therefore, we were interested to see how dysregulation of lipid metabolism could affect ACE2 expression. SREBPs are a family of transcription factors which control lipid synthesis and adipogenesis by controlling enzymes required for cholesterol, fatty acid, triacylglycerol and phospholipid synthesis. In cholesterol deprivation, they translocate from the endoplasmic reticulum to the Golgi apparatus where they are then targeted to the nucleus following cleavage. In the nucleus, they proceed to induce the expression of fatty acid and sterol synthesis (40). SREBP family is composed of SREBP-1 and SREBP-2. SREBP-1 exists as two isoforms: SREBP1-a and SREBP1-c. These isoforms are controlled by independent regulatory proteins and appear to respond differently to different states of lipid factors. Our study suggests that suppressing SREBP1 leads to upregulation of ACE2 expression.

In summary, our findings from the publicly available transcriptomic data show that SARS-CoV-2 infection has significant effects on pathways involved in lipid metabolism. This is of importance as dysregulation of lipid metabolism is a feature of obesity, one of the risk factors of COVID-19. More importantly our data reveal that this increased susceptibility may be due to an increase in ACE2 expression in the lung which is regulated by SREBP1. These findings may potentially aide us in understanding the increased susceptibility in relation to other risk factors such as diabetes and hypercholesteremia.

## References

1. Velavan TP, Meyer CG. The COVID-19 epidemic. Tropical medicine & international health: TM & IH. 2020;25(3):278–80.

2. Lighter J, Phillips M, Hochman S, Sterling S, Johnson D, Francois F, et al. Obesity in patients younger than 60 years is a risk factor for Covid-19 hospital admission. Clinical infectious diseases: an official publication of the Infectious Diseases Society of America. 2020.

3. Patwardhan P. COVID-19: Risk of increase in smoking rates among England’s 6 million smokers and relapse among England’s 11 million ex-smokers. BJGP open. 2020.

4. Li C, Xu MM, Wang K, Adler AJ, Vella AT, Zhou B. Macrophage polarization and meta-inflammation. Translational research: the journal of laboratory and clinical medicine. 2018;191:29–44.

5. Painter SD, Ovsyannikova IG, Poland GA. The weight of obesity on the human immune response to vaccination. Vaccine. 2015;33(36):4422–9.

6. Honce R, Schultz-Cherry S. Impact of Obesity on Influenza A Virus Pathogenesis, Immune Response, and Evolution. Front Immunol. 2019;10:1071-.

7. Hruby A, Hu FB. The Epidemiology of Obesity: A Big Picture. Pharmacoeconomics. 2015;33(7):673–89.

8. Lake MA. What we know so far: COVID-19 current clinical knowledge and research. Clin Med (Lond). 2020;20(2):124–7.

9. Oudit GY, Imai Y, Kuba K, Scholey JW, Penninger JM. The role of ACE2 in pulmonary diseases--relevance for the nephrologist. Nephrology, dialysis, transplantation: official publication of the European Dialysis and Transplant Association - European Renal Association. 2009;24(5):1362–5.

10. Gu H, Xie Z, Li T, Zhang S, Lai C, Zhu P, et al. Angiotensin-converting enzyme 2 inhibits lung injury induced by respiratory syncytial virus. Sci Rep. 2016;6:19840-.

11. Blanco-Melo D, Nilsson-Payant BE, Liu W-C, Møller R, Panis M, Sachs D, et al. SARS-CoV-2 launches a unique transcriptional signature from in vitro, ex vivo, and in vivo systems. bioRxiv. 2020:2020.03.24.004655.

12. Emig D, Salomonis N, Baumbach J, Lengauer T, Conklin BR, Albrecht M. AltAnalyze and DomainGraph: analyzing and visualizing exon expression data. Nucleic Acids Res. 2010;38(Web Server issue):W755–62.

13. Zhou Y, Zhou B, Pache L, Chang M, Khodabakhshi AH, Tanaseichuk O, et al. Metascape provides a biologist-oriented resource for the analysis of systems-level datasets. Nature communications. 2019;10(1):1523.

14. Wu D, Yang XO. TH17 responses in cytokine storm of COVID-19: An emerging target of JAK2 inhibitor Fedratinib. Journal of microbiology, immunology, and infection = Wei mian yu gan ran za zhi. 2020.

15. Stoiber K, Nagło O, Pernpeintner C, Zhang S, Koeberle A, Ulrich M, et al. Targeting de novo lipogenesis as a novel approach in anti-cancer therapy. British Journal of Cancer. 2018;118(1):43–51.

16. Hale AJ, ter Steege E, den Hertog J. Recent advances in understanding the role of protein-tyrosine phosphatases in development and disease. Developmental Biology. 2017;428(2):283–92.

17. Thiriet M. Receptor Tyrosine Phosphatases. In: Thiriet M, editor. Signaling at the Cell Surface in the Circulatory and Ventilatory Systems. New York, NY: Springer New York; 2012. p. 689–703.

18. Shintani T, Higashi S, Suzuki R, Takeuchi Y, Ikaga R, Yamazaki T, et al. PTPRJ Inhibits Leptin Signaling, and Induction of PTPRJ in the Hypothalamus Is a Cause of the Development of Leptin Resistance. Sci Rep. 2017;7(1):11627.

19. Kutmon M, van Iersel MP, Bohler A, Kelder T, Nunes N, Pico AR, et al. PathVisio 3: An Extendable Pathway Analysis Toolbox. PLoS Comput Biol. 2015;11(2):e1004085.

20. Lubis AR, Widia F, Soegondo S, Setiawati A. The role of SOCS-3 protein in leptin resistance and obesity. Acta medica Indonesiana. 2008;40(2):89–95.

21. Marshall R. The Pulmonary Renin-Angiotensin System. Current pharmaceutical design. 2003;9:715–22.

22. Wang J, Chen L, Chen B, Meliton A, Liu SQ, Shi Y, et al. Chronic Activation of the Renin-Angiotensin System Induces Lung Fibrosis. Sci Rep. 2015;5(1):15561.

23. Kickbusch I, Leung G. Response to the emerging novel coronavirus outbreak. BMJ. 2020;368:m406.

24. Vaduganathan M, Vardeny O, Michel T, McMurray JJV, Pfeffer MA, Solomon SD. Renin– Angiotensin–Aldosterone System Inhibitors in Patients with Covid-19. New England Journal of Medicine. 2020.

25. Kuster GM, Pfister O, Burkard T, Zhou Q, Twerenbold R, Haaf P, et al. SARS-CoV2: should inhibitors of the renin–angiotensin system be withdrawn in patients with COVID-19? European Heart Journal. 2020.

26. Patel VB, Basu R, Oudit GY. ACE2/Ang 1-7 axis: A critical regulator of epicardial adipose tissue inflammation and cardiac dysfunction in obesity. Adipocyte. 2016;5(3):306–11.

27. Tilton SC, Waters KM, Karin NJ, Webb-Robertson B-JM, Zangar RC, Lee KM, et al. Diet-induced obesity reprograms the inflammatory response of the murine lung to inhaled endotoxin. Toxicol Appl Pharmacol. 2013;267(2):137–48.

28. Du Y, Guo M, Whitsett JA, Xu Y. ‘LungGENS’: a web-based tool for mapping single-cell gene expression in the developing lung. Thorax. 2015;70(11):1092–4.

29. Plantier L, Besnard V, Xu Y, Ikegami M, Wert SE, Hunt AN, et al. Activation of sterol-response element-binding proteins (SREBP) in alveolar type II cells enhances lipogenesis causing pulmonary lipotoxicity. J Biol Chem. 2012;287(13):10099–114.

30. Shimano H, Sato R. SREBP-regulated lipid metabolism: convergent physiology — divergent pathophysiology. Nature Reviews Endocrinology. 2017;13(12):710–30.

31. Cao X, Yang F, Shi T, Yuan M, Xin Z, Xie R, et al. Angiotensin-converting enzyme 2/angiotensin-(1-7)/Mas axis activates Akt signaling to ameliorate hepatic steatosis. Sci Rep. 2016;6:21592-.

32. Cao X, Lu X-M, Tuo X, Liu J-Y, Zhang Y-C, Song L-N, et al. Angiotensin-converting enzyme 2 regulates endoplasmic reticulum stress and mitochondrial function to preserve skeletal muscle lipid metabolism. Lipids in Health and Disease. 2019;18(1):207.

33. Huang C, Wang Y, Li X, Ren L, Zhao J, Hu Y, et al. Clinical features of patients infected with 2019 novel coronavirus in Wuhan, China. Lancet (London, England). 2020;395(10223):497–506.

34. Chehimi M, Vidal H, Eljaafari A. Pathogenic Role of IL-17-Producing Immune Cells in Obesity, and Related Inflammatory Diseases. J Clin Med. 2017;6(7):68.

35. Hotamisligil GS. Inflammation and metabolic disorders. Nature. 2006;444(7121):860–7.

36. Mackenzie JM, Khromykh AA, Parton RG. Cholesterol manipulation by West Nile virus perturbs the cellular immune response. Cell host & microbe. 2007;2(4):229–39.

37. Zhang J, Lan Y, Sanyal S. Modulation of Lipid Droplet Metabolism-A Potential Target for Therapeutic Intervention in Flaviviridae Infections. Front Microbiol. 2017;8:2286-.

38. Wunderlich CM, Hövelmeyer N, Wunderlich FT. Mechanisms of chronic JAK-STAT3-SOCS3 signaling in obesity. JAKSTAT. 2013;2(2):e23878–e.

39. Jia HP, Look DC, Shi L, Hickey M, Pewe L, Netland J, et al. ACE2 receptor expression and severe acute respiratory syndrome coronavirus infection depend on differentiation of human airway epithelia. J Virol. 2005;79(23):14614–21.

40. Bertolio R, Napoletano F, Mano M, Maurer-Stroh S, Fantuz M, Zannini A, et al. Sterol regulatory element binding protein 1 couples mechanical cues and lipid metabolism. Nature communications. 2019;10(1):1326.

